# BayVarC: an ultra-sensitive ctDNA variant caller using Bayesian approach

**DOI:** 10.1101/2024.02.03.578772

**Authors:** Dongxue Che, Cheng Yan, Jianjun Zha, Zexiong Niu, Quanyu Yang, Xiaolei Cheng, Mipeng Han, Changshi Du, Ke Zhang, Yunfu Hu, Yu-Fei Yang

**Affiliations:** Genetron Health (Beijing) Technology, Co. Ltd., Beijing, China

## Abstract

In liquid biopsy, it is critical to detect variants of allele frequencies as low as 0.1% or even lower, especially when used to monitor secondary resistant mutations and minimal residual disease. Despite the efforts on improving experimental design, it remains challenging to distinguish low-frequency variants from technical noises in the downstream bioinformatic analysis. Here, we introduce BayVarC, a novel variant caller specifically designed for variant calling in liquid biopsy. It applies Bayesian inference to accurately quantify noise level in a locus-specific manner, enabling the discrimination between technical noise and low-frequency cancer variants. Detailed in-silico simulation and in-vitro experiments demonstrated BayVarC’ superior performance over existing state-of-the-art tools. BayVarC can effectively detect low frequency variants while maintaining low false positive rate (0.05 FP/KB). Meanwhile, it achieves Limit of Detection (LoD) as low as 0.1%. Furthermore, empowered by its architecture, BayVarC shows promising applicability in Minimal Residual Disease (MRD) detection. BayVarC is freely available at https://github.com/GenetronBioinfomatics/BayVarC.

## Introduction

A decade ago, the use of High Throughput Sequencing (HTS) technology in clinical oncology was limited and primarily used to detect medically actionable cancer variants in tissue biopsy^1–4^. As HTS technology progresses, it has unlocked numerous new applications in clinical oncology^5–9^. For instance, nowadays it has been widely used in liquid biopsy: to assess the efficacy of cancer drugs before administration (companion diagnostic)^10, 11^; to monitor the sign of Minimum Residual Disease (MRD) in postoperative care^12–14^; to detect early cancer signal in individual of high cancer risk^15–17^. Liquid biopsy owns numerous advantages over tissue biopsy, such as being non-invasive and overcoming tumor heterogeneity^18–20^. In the meantime, these applications have brought new challenges. In tissue biopsy, moderate Limit of Detection (LoD) of variant detection (∼1%) is commonly considered to be sufficient^21^. However, liquid biopsy requires detection of low frequency variant since, in most instances, only a small proportion of tumor DNA can be elicited to circulation system^22, 23^. To apply liquid biopsy in companion diagnostic, we usually expect to detect variants of frequency below 1%^10^. For MRD and early cancer detection, mutations are expected to be detectable at frequency as low as 0.02% or even lower^24, 25^. The dilemma is that many steps of HTS experimental procedures, such as sample storage, PCR amplification and sequencing, may generate false positive variants (technical noise) and most of them are of low frequency. Although some efforts have taken at experimental step, such as the introduction of Unique Molecular Identifiers (UMIs), it only mitigated the problem. More works are in need to better handle the technical noise before HTS based liquid biopsy can provide more accurate guidance in clinical practice.

Variant calling algorithms have been actively investigated along with the development of HTS technology, yielding many publicly available variant callers. Roughly speaking, they can be categorized by purpose (SNV only vs SNV/ InDel), by prerequisite (tumor only vs tumor-normal paired) or by algorithm. For instance, Lofreq, a frequently used low frequency variant caller, can call SNV as well as InDel in tumor only mode. It applies allele frequency analysis approach which directly models allele frequency to detect cancer variants^26^. This approach is considered as one of the main algorithms suited for low frequency variant detection^27–29^ and has been adopted by many other variant callers, for instance, MuTect ^30^ and deepSNV^31^. Another state-of-the-art low frequency variant caller, VarDict, calls SNV, InDel and SV simultaneously in tumor-only mode. It uses heuristic approach to perform variant calling^32^. The same approach has also been applied in VarScan2^33^. Some other callers, such as Pisces^34^ and SNVer^35^, estimate error levels and/or distinguish cancer variant based on specific distributions. Despite that some algorithms are believed to be suited for calling low frequency variant, we found that they are not sufficiently sensitive to call variant of frequency lower than 1% while remain low false positive variants rate. Additionally, in some studies^36, 37^, hard filtering approaches are still being used on ctDNA variant detection. These approaches set unified thresholds of variant frequency and count of variant supporting reads. However. The technical noise on each genomic locus is not the same due to some intrinsic genomic characteristic. Among some genomic loci, the differences in noise level may be as large as two orders of magnitude. This presents a drawback for setting unified thresholds in variant detection.

Here we introduce BayVarC, a variant caller tailored for detecting low frequency variants in liquid biopsy. It achieves high sensitivity on low frequency variants by constructing locus-specific error model. During the error modeling, We applied a Bayesian inference which allows us to train BayVarC model using limited samples while ensure accurate estimation of locus-specific error. BayVarC can detect SNV and InDel simultaneously in tumor-only mode. Additionally. we developed InDel signal booster inside BayVarC to further improve its detection of low frequency InDel. We conducted comprehensive benchmarking using in-silico simulation, public in-vitro data and commercial reference standard samples. Results showed that BayVarC outperformed currently available cutting-edge methods and can be reliably used in liquid biopsy.

## Material and Methods

### Feature engineering and modeling

The SNV and InDel model in BayVarC was trained separately using a dataset of 40 non-cancerous samples (see **Data generation** for details). Briefly, at each genomic locus *p*, the observed counts of background alternative allele (allele that differ from reference genome hg19 at given genomic locus) *y*_*p*_ under the given total coverage *N*_*p*_ has a locus-specific baseline noise rate *x*_*p*_. The model was then assumed to be fitted with a Binomial distribution as:

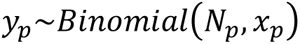

If sample size is sufficiently large, we can set the observed mean error rate of given genomic locus as parameter. However, with insufficient samples, we introduced Bayesian inference to borrow information from genomic loci with same genomic features. Feature types used in SNV model include tri-nucleotide context and mappability. Mappability is a binary feature, indicating whether the genomic position was uniquely mapped. Each genomic locus was assigned to corresponding group based on its feature. Error frequencies for each group were assumed to be drawn from a Beta distribution, the same feature group were fitted with same parameters. A tri-nucleotide context group *i* (*i* = 1, …, 64) and mappability information *j* (*j* ∈ {*unique alignment*, *multiple alignment*}) at genomic locus *p* (*p* = 1, …, *P*) were combined to fit Beta distribution:

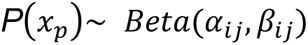

For genomic locus *p* in the same feature group, Beta distribution was applied to estimate the group shared prior error rate using the method of moments. For given genomic locus *p*, the mean observed error rate in 40 non-cancerous samples was defined as likelihood. its posterior error rate follows:

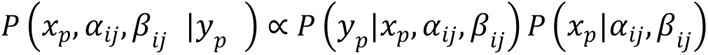

Different from SNV, features in InDel model include segmental duplication, DNA sequence complexity (non-repeats, simple repeat, LINE, SINE, LTR, DNA and low complexity sequence) and InDel length. Similar to SNV model, genomic loci were grouped based on their features. For a genomic locus *p* (*p* = 1, …, *P*), segmental duplication information group *i* (*i* ∈ {*SegDup*, *NonSegDup*}) and DNA sequence complexity group *j* (*j* ∈ {*LINE*, *SINE*, *LTR*, *DNA*, *Simple repeat*, *Low complexity*, *Nonrepeats*}) were combined to fit model. Group shared prior error rates were estimated. Locus-specific posterior error rate was then estimated based on Bayesian theorem.

Based on the procedure described above, we were able to constructed locus-specific error model. In practice, we compare observed signal in ctDNA samples with model specified posterior error rate. Binomial distribution with model defined p-value cut-off is performed to determine whether the observed signal represents true variant or technical noise.

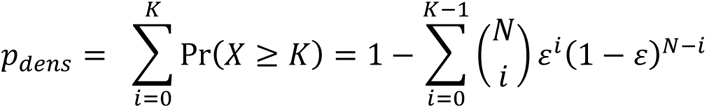

Where *N* is sequencing depth of given locus, *K* is alternative allele count and *ε* is posterior error rate specified by our model.

### Sample processing

Same wet-lab procedure was applied to process cfDNA samples used in BayVarC model construction as well as ctDNA samples used in a series of validations. DNA was extracted from the plasma following the manufacturer’s protocol by the MagMax^TM^ Cell-Free DNA Isolation Kit (Thermo fisher A29319). DNA Fragment size was evaluated using 4200 TapStation (Agilent Technologies). cfDNA was quantified by a fluorescent assay using the Qubit 4.0 Fluorometer (Thermo Scientific, Waltham, USA). Sequencing libraries were prepared using the IDT xGen Prism Library Kit (Integrated DNA Technologies, Inc, (IDT)). Unique molecular identifiers (UMI) were incorporated in a unique, single-stranded ligation strategy. cfDNA of 172 tumor-related genes was captured using a commercial assay Onco Sonar^TM^. The enriched libraries were amplified with P5/P7primer. After qualified by the 4200 TapStation and quantified with Qbit4.0 and a qPCR NGS library quantification kit (Agilent Technologies), Libraries were sequenced on a NovaSeq 6000 platform (Illumina, San Diego, CA) or MGISEQ-2000 platform (MGI Tech Co., Ltd).

### Pre-processing of raw sequencing reads

8-bp UMI sequences were trimmed from raw sequence data (in FASTQ format) and then added to the sequence header and adapter sequences were trimmed using Trimmomatic (version 0.33) with parameters “LEADING:3 TRAILING:3 SLIDINGWINDOW:4:15 MINLEN:36”. Paired-end clean reads were first mapped to the human reference genome (hg19) using the Burrows-Wheeler Aligner (BWA, version 0.7.10-r789) by default parameters. Then consensus sequence reads were created based on UMI, mapping positions and insert sizes. Consensus FASTQ files were re-aligned to the human reference genome (hg19) using the Burrows-Wheeler Aligner. mpileup (v1.7) was used to count initial sequencing depth of reference allele and alternative allele at each genomic locus.

### InDel booster

To rescue reads that contain InDel signals, but are mislabeled as softclips by alignment software, we developed a model named ‘InDel booster’. For each InDel candidate (genomic locus with alternative DNA sequence), InDel booster retrieves surrounding softclip reads and performs BLAST-like local realignment within 100bp up- and down-stream of InDel signal. Based on the realignment results, InDel booster infers the nature of softclips and then re-calculate the frequency of InDel signal. Through this frequency recalibration process, some InDels may be identified due to elevated allele frequency.

### In-silico validation

Our aim is to evaluate the detection accuracy of BayVarC using ctDNA samples with labeled variants of known allele frequency. To do this, we used 25 cfDNA samples previously collected (out of samples used in model building). BAM files were generated (previously described procedure) and used as background templates. To make the evaluation more realistic, we collected variant information from The Cancer Genome Atlas (TCGA) project and generate a variant sampling pool. Briefly, MAF files of five cancer types (BRCA、LUAD、STAD、LIHC and COAD) were retrieved from GDC Data Protal (https://portal.gdc.cancer.gov/). For each cancer type, MAF files were merged into one single variant pool. BAMSurgeon (version 1.4.1) was used to randomly sample from variant pool and insert variants into BAM files alone with their original allele frequency. Finally, five BAM files were generated for each cancer type and each BAM file contains 50 SNVs and 50 InDels. Variants in each BAM file were then randomly and equally split into five portion and pre-specified their allele frequency as 0.1%, 0.25%, 0.3%, 0.5% and 1% BAM files were then subject to variant detection by BayVarC and other variant callers, including VarDict, LoFreq, VarScan2, MuTect, Pisces and SNVer. As some studies utilize a unified hard filtering approach with a fixed mutant allele cutoff, we included this strategy in our comparison, referred to hereafter as ‘Fixed_parameters’.

### In-vitro validation

In-vitro validation was carried out to objectively evaluate the performance of BayVarC. We acquired Twist cfDNA Pan-cancer reference standard samples from Twist Bioscience (San Francisco, CA, USA) and Seraseq® ctDNA Complete™ Mutation Mix standard samples from SeraCare (Milford, MA, US). Samples were then sequenced and pre-processed through experimental procedure described above, and then analyzed by BayVarC to estimate both sensitivity and limit of detection (LoD).

In addition, we retrieved in-vitro ctDNA FASTQ data generated by Burning Roch.Inc in FDA-led Sequence Quality Control 2 Project (SEQC2)^36^. Experimental workflow to produce the data was described in the study. Retrieved FASTQ files were then processed and variants were called by each callers including BayVarC.

### Simulation of MRD detection using BayVarC

Twist cfDNA Pan-cancer reference standard samples of variant allele frequency 0.2% were used for simulation. To meet MRD criteria, cfDNA samples were subjected to in-silicon dilution to obtain variants with an allele frequency of 0.02%. 20 variants were selected randomly to simulate an MRD-positive sample, and the same 20 variants were tracked in blank samples to simulate an MRD-negative sample. 10,000 in-silico simulations were performed. Samples with two or more detected variants were considered as MRD positive. Performance was then evaluated through parameter space.

## Results

### Overview of BayVarC pipeline

BayVarC employs a Bayesian inference to construct a locus-specific noise model using 40 non-cancerous cfDNA samples (See Methods). In practice, BayVarC compares the observed signal, indicative of candidate variant, against the posterior error rate as defined by the BayVarc model. Subsequently, BayVarC employs Binomial testing with predefined significance level (1*10^5) to determine the nature of observed signal (Fig 1A and Supplementary Fig. 1).

**Figure 1.**
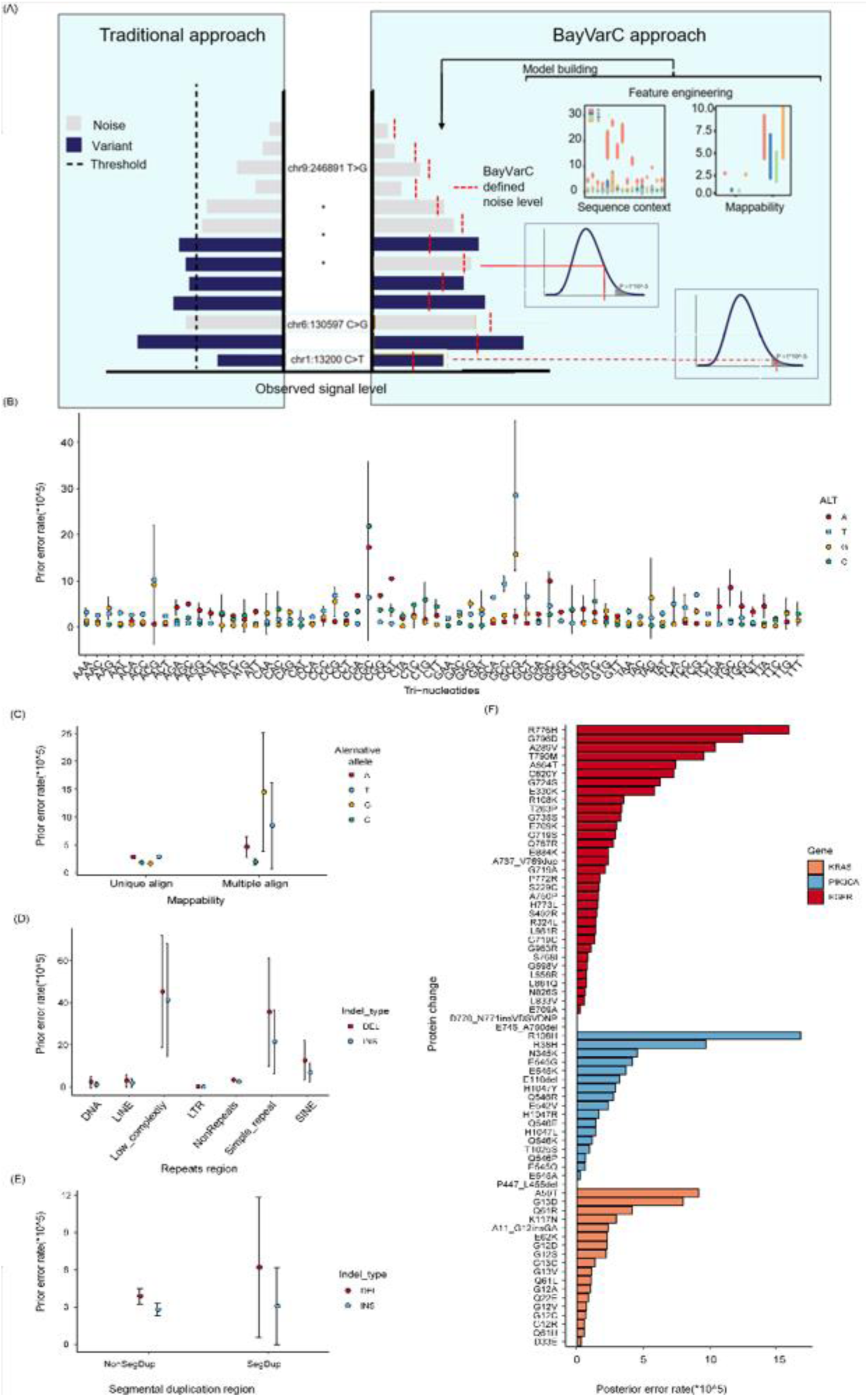
The BayVarC model. (A)Workflow of BayVarC model. Compared with tranditional approach (left panel), BayVarC (right penal) utilizes Bayesian approach to construct locus-specific error rate (vertical dash line) which is further used to compare observed signal level (horizontal grey / blue bar) under Binomial distribution to determine the nature of observed signal. (B)Prior error level with 95% confidence interval across different trinucleotide context in SNV model. (C)Prior error level in uniquely aligned region and multiple alignment region in SNV model. (D)Prior error level across different repeat regions in InDel model. (E)Prior error level in Segmental duplication region and non-segmental duplication region. (F)Posterior error level of oncogenic hotspots in EGFR, KRAS and PIK3CA.

### The BayVarC model

The primary challenge in developing ctDNA error model is the precise estimation of locus-specific background error using limited samples. BayVarC addresses this challenge by implementing a Bayesian inference, wherein prior knowledge is incorporated to enhance the accuracy of error estimation. However, the accuracy of prior knowledge depends on the selection of appropriate genomic features. In the SNV model, features such as tri-nucleotide sequence context and mappability are considered, while the InDel model incorporates sequence complexity elements (See Method). Figure 1B-E showed the distribution of prior error rate across different feature group in both SNV and InDel models. In SNV model, some tri-nucleotide sequence context, such as ACG, GCG and CGC, are significantly more error prone than others (Fig 1B). The error rate is 10 folds greater compared with most of other tri-nucleotide sequence context. This observation coincides with previous studies^38–41^. Furthermore, genomic regions of multiple mapping are more error prone than uniquely mapped region (Fig 1C). In InDel model, the error rate varies significantly between different repeat region as well as different segmental duplication region (Fig 1D-E). As expected, low complexity region, highly repeat region and segmental duplication region showed elevated error rate. The elevated error rate in those regions probably due to sequence misalignment and/or dephasing during sequencing ^42^. In both models, the variance in prior error rates among feature groups rationalizes our feature engineering approach, demonstrating that error rates of genomic group provide essential prior knowledge for accurate posterior error estimation.

The posterior error rate of each genomic locus was calculated based on Bayesian theorem (See Method). Across the panel, the median SNV and InDel posterior error rate is 1.67e-05 and 2.38e-05 respectively. However, the posterior error rate varies significantly between genomic loci. Figure 1F showed posterior error rates of some well-known oncogenic hotspots. For instance, EGFR T790M owns much higher error rate than EGFR L858R (9.58e-05 vs 7.34e-06), which is consistent with earlier observations. With this locus-specific error model, we were able to set cutoff in a locus-specific manner.

Another factor that affects accuracy of the model is the number of cfDNA samples used to estimate the background error rate. Insufficient sample size during model training may lead to biased posterior error rate at some genomic loci. We therefore estimated the appropriate sample size for generating stable model. As showed in Supplementary Fig 2A, In SNV model, the error level of most context denoted curves reached stable state when simple size approached to 40. In InDel model, error level of most feature groups remained low. Notably, Feature group of ‘low complexity’ exhibited highest error level and stabilized when sample size approached to 40 as well (Supplementary Fig 2B). In both SNV and InDel models, the R^2^ (coefficient of determination) between noise calculated using different sample size reached to 1 when sample size approached to 40 (Supplementary Fig 2C).

### Performance comparison using in-silico data

We then assessed the performance of BayVarC against other well-known variant callers using 25 cfDNA in-silico samples in which synthetic variants (allele frequencies ranging from 0.1% to 1%) were inserted (See Method). Through random selection from TCGA project, a total number of 834 SNVs and 1091 InDels were successfully inserted, while 416 SNVs and 119 InDels were failed to meet BamSurgeon’s insertion criteria and hence were unsuccessfully inserted (Supplementary Table 1). Our initial comparison included established variant caller such as VarDict, LoFreq, VarScan2, MuTect, Pisces, SNVer and fixed parameter. However, we found that MuTect, Pisces and SNVer can only detect few variants of frequency lower than 1%. Consequently, we concluded that these three variant callers are not suitable for ctDNA variant calling and therefore excluded them from further benchmarking. We classified all detected variants, except those in-silico inserted variants, as false positives if they were not germline variants with frequency >= 20%^43^. The Area Under the Receiver Operating Characteristic (AUROC) curve revealed that BayVarC significantly outperformed others in both SNV and InDel detection (Fig 2A-B). The parameters of each caller used in the comparison and corresponding performance were detailed in Supplementary Table 2.

**Figure 2.**
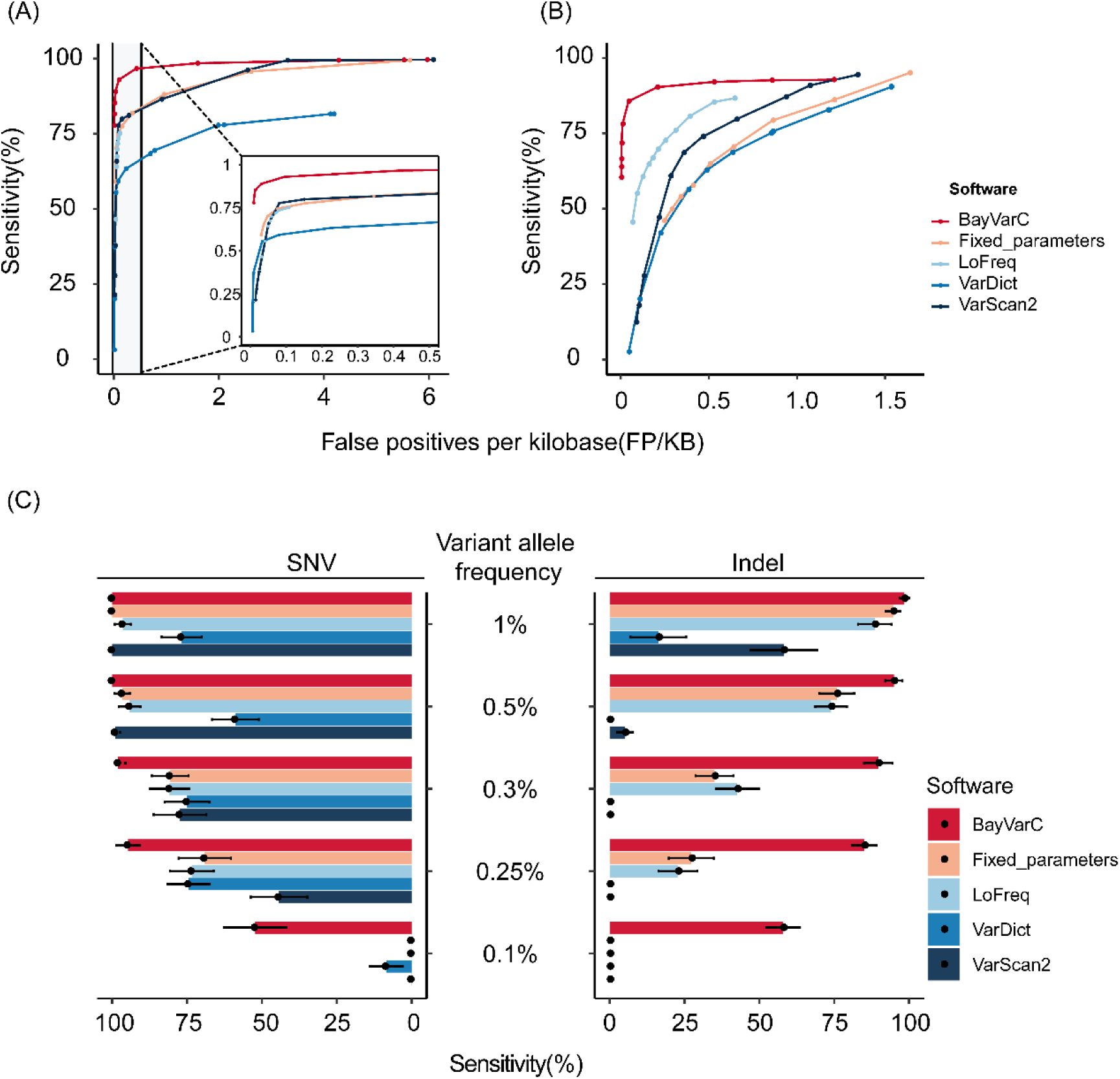
AUROC comparison between variant callers using in-silico simulation data. (A)Performance of SNVs. Left side was shown in addition zoom in window. (B)Performance of InDels. (C)Sensitivity comparison when FPR is fixed at 0.05 FP/Kb

We first fixed the false positive rate (FPR) at 0.05 false positive per kilobase (FP/KB) for all callers, as per average FPR in SEQC2 study^36^. With such specificity setting, most callers perform poorly while BayVarC can still detect averagely over 85% of SNV and InDels variants across allele frequency range (Fig 2C and Supplementary Table 3). Especially for low-frequency variants (0.1%-0.3%), BayVarC exhibits significantly higher sensitivity than other variant callers. To explore the performance in scenarios where a slightly higher FPR is accepted, we relaxed FPR to 0.5 FP/KB. Results showed that BayVarC continued to outperform other variant callers in terms of detection sensitivity, especially for low-frequency variants (Supplementary Table 4). Conversely, when we fixed sensitivity at 0.75 for all callers, most variant callers yielded significantly more artifacts compared to BayVarC (Fig 3 and Supplementary Table 5). Upon closer examination, a significant proportion of artifacts detected by these tools were associated with high noise levels, e.g., C>T substitutions. Notably, BayVarC enhances the signal strength required for detecting variants at these sites, thereby effectively eliminating these false positive variants. Altogether, these observations underscored the superior performance of BayVarC, attributed to its well-crafted feature engineering process as well as its Bayesian architecture.

**Figure 3.**
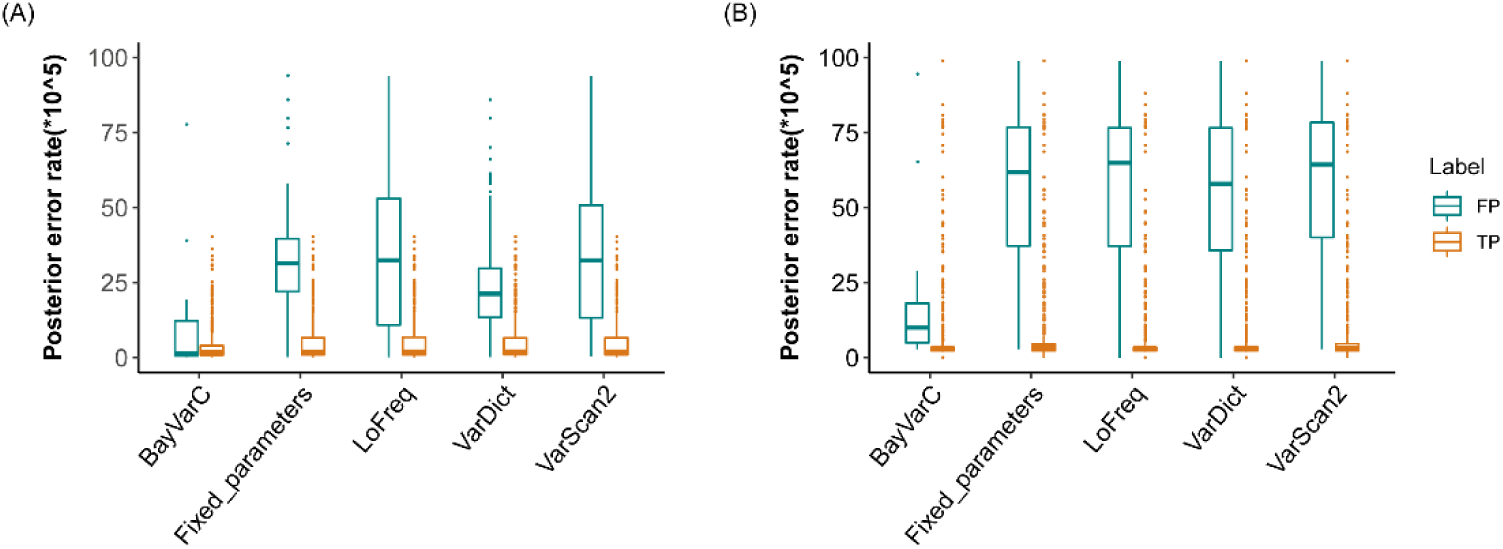
The error rate distribution of detected variants when sensitivity was fixed at 0.75 within in-silico data. Different colors indicate true positive detection (TP) and false positive detection (FP). (A) Box plot of error rate of SNVs detected by variant callers. (B) Box plot of error rate of InDels detected by variant callers.

### Performance comparison using in-vitro data

To ensure a more objective assessment of performance, we further conducted an in-vitro validation using commercial reference standard ctDNA samples (See Methods and Supplementary Table 6). The reason we choose reference standard samples is that they provide positive variants with quantified allele frequency, facilitating both sensitivity and limit of detection (LoD) evaluation. To enhance the representativeness of the results, we selected reference standards with different characteristics from both Seraseq® ctDNA Complete™ Mutation standard and Twist cfDNA Pan-cancer reference standard for testing. Each reference standard underwent testing 12 or 20 times, with a total number of 133 labeled SNVs and 139 labeled InDels for evaluation. For clarity of comparison, we fixed DNA input as 30ng and sequencing depth as 28,000X. To ensure comparability as well as high specificity required by liquid biopsy, parameter values of all callers were set to achieve FPR close to 0.05 FP/KB, referring to the results of in-silico simulations. Results showed that BayVarC consistently achieved significantly higher sensitivity across allele frequency ranges, particularly in low allele frequency groups (Fig 4A). Notably, BayVarC exhibited superior performance in reaching lower LoD for both SNVs and InDels; for instance, 7/24 SNVs and 4/12 InDels reached a LoD of 0.125%. Furthermore, even though the sensitivities of detecting SNVs with allele frequency of 0.5% were comparable among all tools (Fig 4A), BayVarC displayed improved detection stability (Fig 4B). Enhanced sensitivity of BayVarC remained when parameter values were adjusted to achieve a higher FPR of 0.5 FP/KB (Supplementary Fig 3), and this conclusion held true for both MGI and illumina sequencing platform (Supplementary Fig 4).

**Figure 4.**
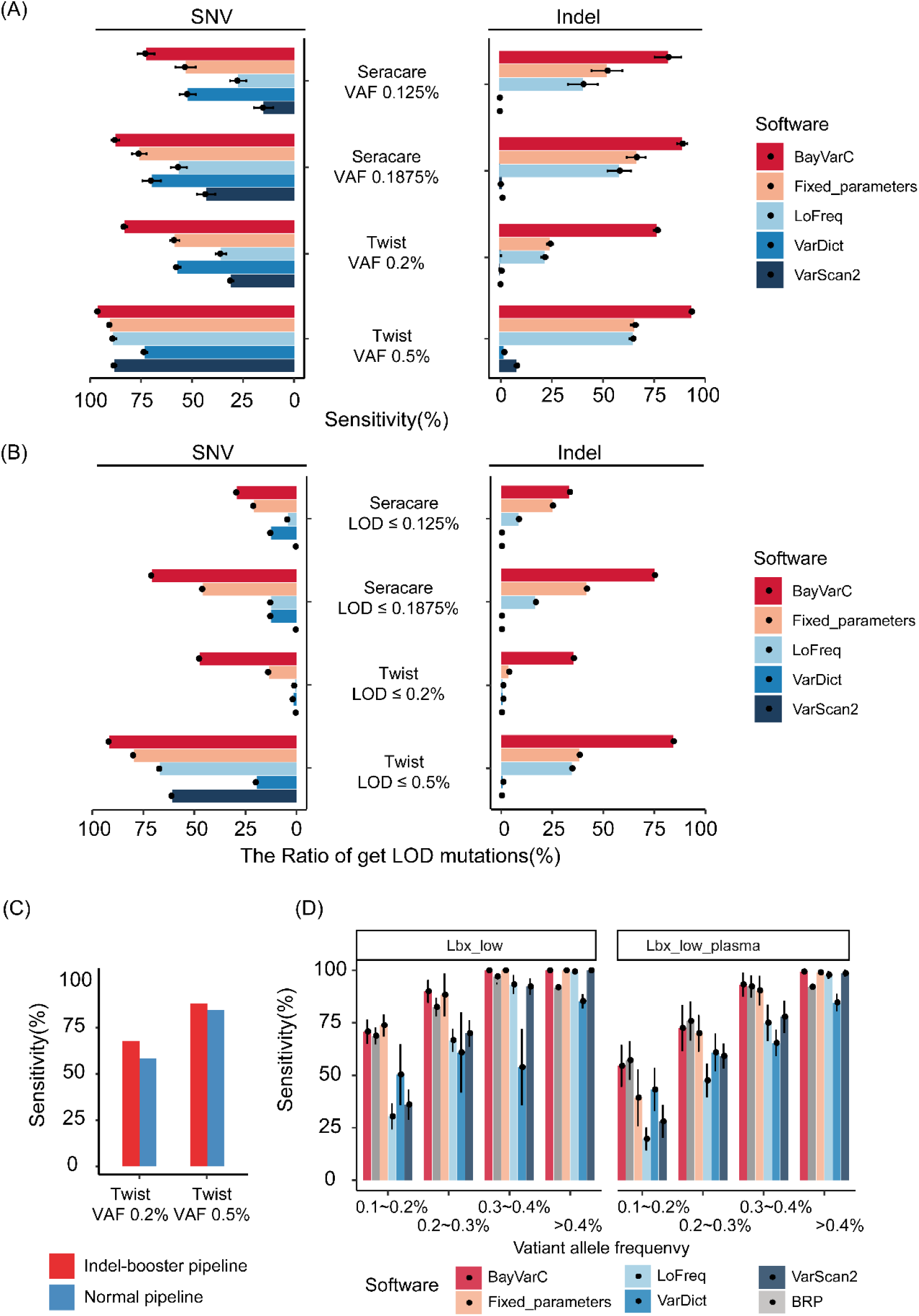
Performance comparison across variant callers using in-vitro data generated from S2000 sequencing platform. (A) Sensitivity comparison across variant callers. (B) Proportion of genomic loci reaching LOD by different variant callers. (C) Sensitivity of InDel booster. (D) Sensitivity comparison on SEQC2 data. Samples in Lbx_low panel were prepared from cfDNA while samples in Lbx_low_plasma were prepared from plasma.

Alignment softwares, such as BWA, often encounter challenge where InDel signals near the ends of the reads are mistakenly labeled as softclips. As the consequence, such InDel variants, especially of low frequency, may be missed by variant caller. To address this issue, we developed ‘InDel booster’ (See Method). Twist cfDNA Pan-cancer reference standard samples contain 28 long InDels and therefore was used to evaluate the performance of InDel Booster. As showed in Figure 4C, When InDel booster was enabled, BayVarC was able to detect 9% more long InDels in Twist cfDNA Pan-cancer reference standard (VAF 0.2%). Furthermore, it achieves 250% increase in the number of long InDels reaching LoD of 0.2% (7 vs 2), demonstrating a significant enhancement in detection stability of long InDels.

Moreover, we thought to assess BayVarC’s performance using publicly available data. The data generated by SEQC2 study was selected for benchmark owing to its study characteristics. To ensure comparability, only FASTQ files started with 25ng DNA were used for benchmark because some callers / parameters are sensitive to DNA input. Results in Figure 4D showed that, in detecting variants of low allele frequency, BayVarC outperforms most of other variant callers including Burning Rock whose performance was directly derived from SEQC2 publication^36^. These benchmark samples were prepared by Sequencing Quality Control (MAQC) consortium and experimentally processed outside of our facility. We demonstrated that, despite the difference in sample preparation procedure, BayVarC exhibited outstanding performance and extended adoptability.

### Potential usage in MRD detection

BayVarC provides a straightforward way to control FPR-sensitivity balance by adjusting its primary parameter, the probability of Type I error (alpha). This adaptability renders BayVarC suitable for a wide range of applications, from broad companion diagnostics demanding high detection specificity to the monitoring of Minimal Residual Disease (MRD) where, within some approaches, moderately decreased mutation-wide specificity is acceptable.

Using diluted Twist cfDNA Pan-cancer reference standard samples (See Method), We investigated the impact of relaxing alpha value on detection sensitivity as well as specificity. As expected, Figure 5A showed a notable linear correlation between alpha value and sensitivity. For instance, at a sequencing depth of 28,000X, the sensitivity for detecting variants with a frequency of 0.1% was increased by 69% when alpha was relaxed from 10^-5^ to 10^-2^. On the other hand, relaxed alpha adversely correlates with FPR across different sequencing depth (Fig 5B). Generally, the value of alpha can be considered as an approximate estimate of FPR. For tumor-informed MRD detection, it is acceptable to moderately increase FPR for individual variant detection, primarily for two reasons: (1) usually only a few variants are tracked, and (2) typically, a sample is considered as MRD positive only when two or more variants are detected^44–46^. Therefore, we attempted to set a relaxed alpha to further enhance the detection of extremely low-frequency variants.

**Figure 5.**
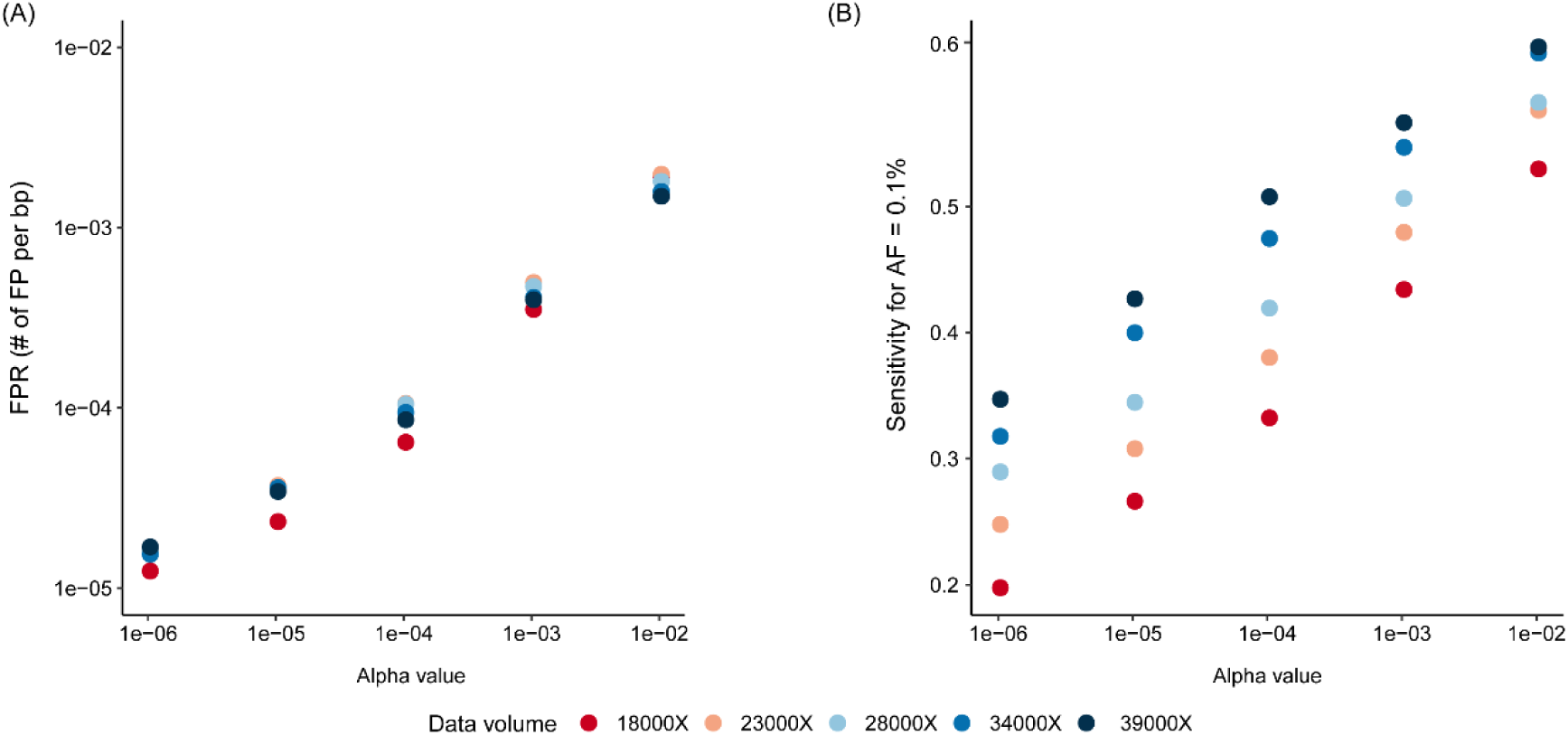
Sensitivity – FPR balance within parameter space of BayVarC across different data volume. (A) The correlation between FPR and alpha. (B) The correlation between sensitivity and alpha.

Based on these observations, we explored the application of BayVarC in MRD detection. We hypothesized a scenario in which a sample is classified as MRD-positive if two or more variants are detected among 20 variants being tracked^44–46^. A FPR of as low as 1% for MRD-negative samples being mis-classified as MRD-positive is acceptable for MRD detection^47^. Setting alpha to 0.01 can approximate this FPR, as inferred from binomial distribution. To confirm the alpha setting as well as detection performance, we ran BayVarC on a set of simulation MRD data (See Method) with different alpha values. As we expected, setting alpha to 0.01 enabled the reliable detection of MRD-positive samples with a positive predictive agreement (PPA) of at least 95%, while maintaining a low false positive rate, as shown in Figure 6.

**Figure 6.**
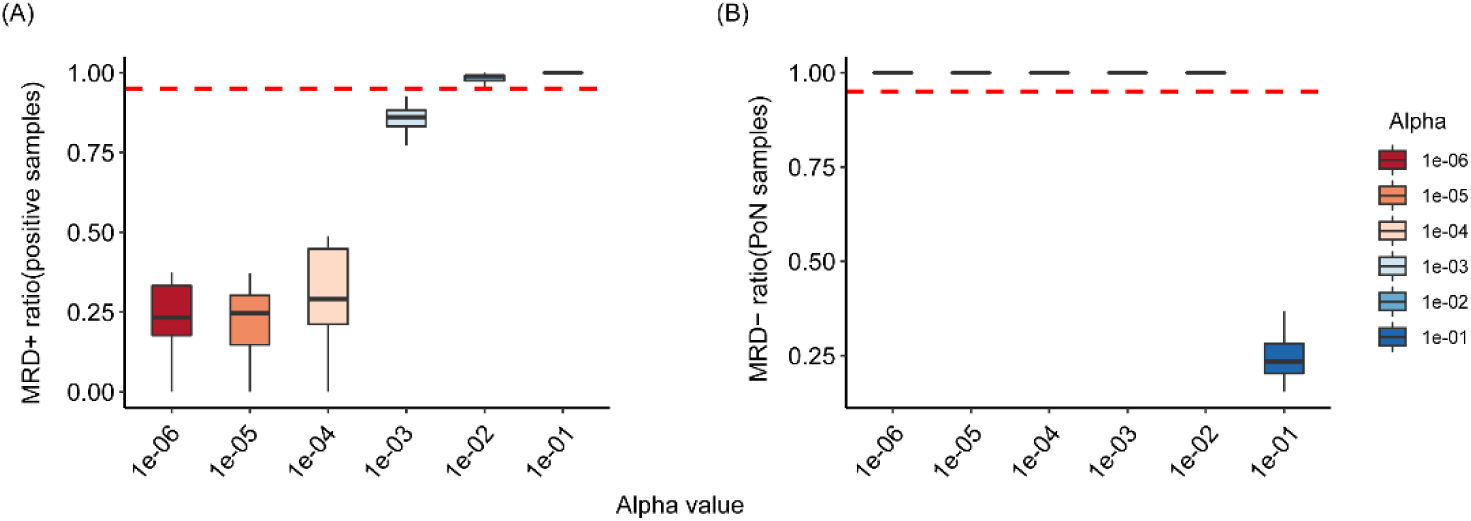
The performance on MRD detection. Dashed horizontal line indicates ratio of 0.95. (A) Box plot of MRD positive rate with different alpha. (B) Box plot of MRD false positive rate with different alpha.

## Discussion

Traditionally, two approaches have been widely used to improve detection of low frequency variants: (1) amplifying the variant signal, for instance, by enriching mutant DNA strand or removing wild-type DNA strand^48, 49^. (2) reducing background noise, such as integrating molecular barcodes during library preparation^50^. Compared to these approaches, BayVarC is designed to accurately quantify background noise and it achieves this goal solely through bioinformatic algorithmic innovation. This approach renders it complementary to other experimental techniques, potentially further enhancing detection efficacy.

Due to the widespread use of error suppression techniques, such as unique molecular identifiers, in liquid biopsies, the technical noise level typically ranges from 10^-6^ to 10^-4^. To precisely measure technical noise of such level, over 1.5 million DNA molecules are required (margin of error = 5*10^-6^, Confidence level = 95% and we use Binomial distribution to approximate the process of high throughput sequencing). This corresponds to over 400 non-cancerous individuals, each providing 10 ml of peripheral blood. In a previous study, up to 1000 negative samples are in need to establish the noise model during MRD testing^44^. This is economically impractical in most scenarios.

In BayVarC, we introduced Bayesian inference. The main idea is to explore genomic features that influence noise level and assume that genomic locus with same features share similar noise level. Hence, genomic loci in the same feature group can share information with each other and use the same prior knowledge in the Bayesian framework. This strategy allowed us to precisely estimate the locus-specific error rate using as few as 40 training cfDNA samples.

Following the establishment of the locus-specific background noise, we can fine-tune the balance between sensitivity and specificity by setting the alpha value of hypothesis testing. For genomic loci exhibiting significant differences in noise levels, a unified alpha value enables differentiation in setting the cutoff for the number of mutant alleles. Consequently, in comparison to the conventional approach of employing a unified hard filtering with a fixed mutant allele cutoff, this strategy of setting variant-specific cutoff optimizes assay performance, by minimizing the false positive results in high-noise loci and maximizing sensitivity in low noise loci.

In the performance comparison, we demonstrated that BayVarC significantly outperformed other variant callers. Further investigation revealed that improved accuracy is mainly reflected from error prone genomic loci, such as error prone trinucleotide context. We understand that most variant callers used for comparison were not originally designed for liquid biopsy and may not be able to perform well on variants of low frequency. Thus, hard filtering based on unified threshold has been used in many ctDNA analysis pipeline. However, unified threshold is not the optimal solution due to distinct error level across genomic loci and, expectedly, showed inferior performance than BayVarC.

Another advantage of BayVarC is that, once the model is constructed, we can forecast the minimum sequencing depth required for achieving desired sensitivity or LOD. Whether in oncology research or diagnostic, some genes/genomic loci are more significant than others. With BayVarC model, users can tailor sequencing depth in a way that genomic locus of interest, for instance cancer related hotspots, can reach desired sensitivity/LoD while less relevant genomic regions are designed in a cost-effective manner.

Admittedly, BayVarC was modeled on limited features. Although we have proved these features to be informative, other factors may influence technical noise. Such factors may include sequencing platform, reagents and batch effect. Therefore, we extended our validation to include public data from SEQC2 project. the primary goal of SEQC2 project is to develop reference materials that suit for evaluation of impact of experimental and bioinformatics variables on variant detection as well as the assessment of inter- or intra-lab reproducibility. The meticulous sample preparation and independent experimental settings of SEQC2 render its data an ideal benchmark for assessing BayVarC’s adaptability. Our findings indicate that BayVarC maintained its sensitivity when applied to SEQC2 data.

Last but not least, we recognize that experimental environment varies between laboratories and our evaluation can not be exhaustive. To fully exploit BayVarC’s strength, we encourage users to train BayVarC error model based on their own experimental environment.

In this study, we demonstrated that BayVarC represents an ultra-sensitive variant caller powered by its Bayesian architecture as well as proper feature engineering strategy. The superior performance in detecting variants of low frequencies makes BayVarC suitable in liquid biopsy where main challenge is to distinguish true variants of low frequencies from technical noise. As liquid biopsy rapidly extends its utility in both oncology research as well as clinical oncology, we believe that BayVarC can facilitate oncology research as well as accelerate the clinical translation of oncology.

## Data availability

BAM files used in in-silico validation are available at NCBI Sequence Read Archive (SRA). FASTQ files of Twist cfDNA Pan-cancer reference standard samples are available NCBI Sequence Read Archive (SRA). FASTQ files of Seraseq® ctDNA Complete™ Mutation Mix standard samples are available NCBI Sequence Read Archive (SRA).

## Code availability

Source code and executables of BayVarC model is freely available at https://github.com/GenetronBioinfomatics/BayVarC. The codes to reproduce the analysis results and generate figures in this manuscript are stored permanently at Figshare.

## Acknowledgement

We would like to thank Yang Du and Zhen Li for their valuable suggestions during the proofreading of manuscript.

## Contributions

Cheng Yan and Yu-Fei Yang designed and initiated the study. Dongxue Che developed the algorithm and conducted the validation. Jianjun Cha and Zexiong Niu helped with data analysis. Quanyu Yang, Xiaolei Cheng, Mipeng Han, Changshi Du and Ke Zhang conducted wet-lab experiments. Cheng Yan wrote and edited the manuscript with input from other authors. Dongxue Che worked on data visualization. Cheng Yan, Yu-Fei Yang and Yunfu Hu led the project.

## Ethics declarations

