## Supplementary Figures 1-4 for "BayVarC: an ultra-sensitive ctDNA variant caller using Bayesian approach"

**Supplementary Figure**

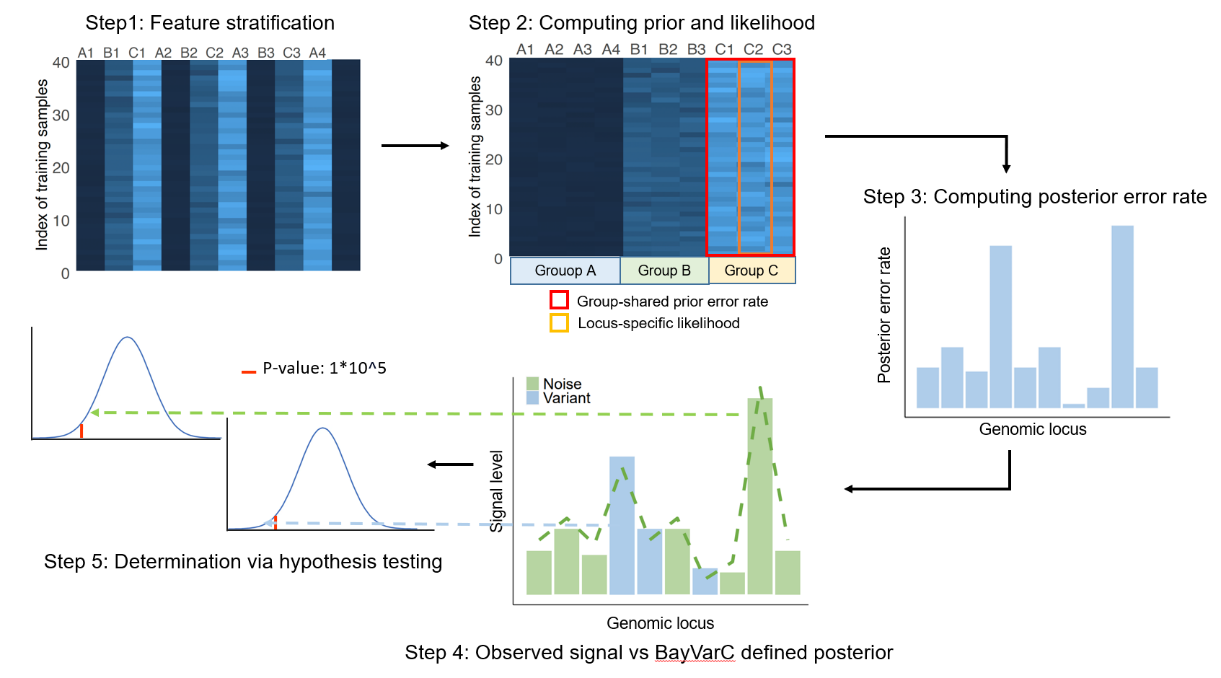

**Supplementary figure 1:** The workflow of BayVarC model. Step 1: Genomic loci of 40 non-cancerous cfDNA samples were stratified based on pre-defined genomic features. Step 2: Group-shared prior error rate and locus-specific likelihood were computed. Step 3: Posterior error rate was computed using Bayesian approach. Step 4: Observed signal level (green / blue bar) at each genomic locus was compared to BayVarC defined posterior error rate (green dash line). Step 5: Hypothesis testing was used to determine whether the observed signal represents technical noise or true variant.

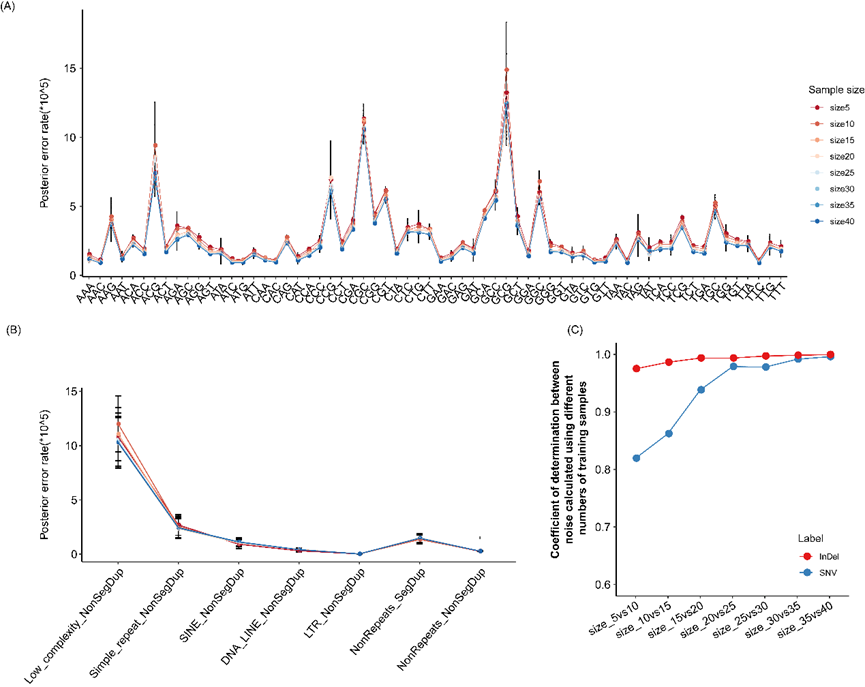

**Supplementary figure 2:** Correlation between noise level and different numbers of cfDNA training samples. (A) The error rate distribution across different tri-nucleotide sequence context using different sample sizes in SNV model. (B) The error rate distribution across different repeat and segmental duplication region using different sample sizes in InDel model. (C) Coefficient of determination between noise calculated using different number of training samples.

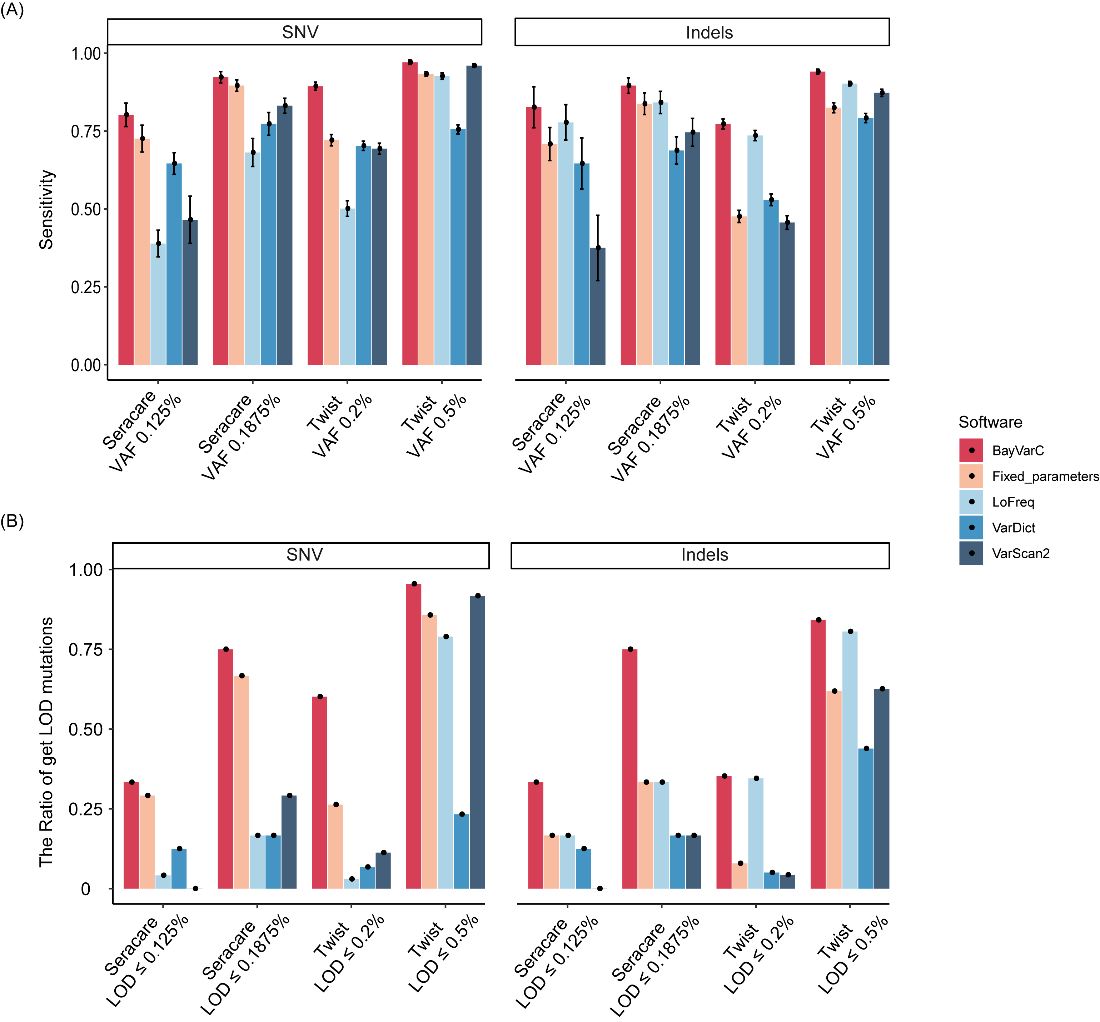

**Supplementary Figure 3:** In-vitro performance of each variant callers when FPR was fixed at 0.5 PF/Kb on S2000 platform. (A) Sensitivity benchmark across variant callers. (B) LoD benchmark across variant callers.

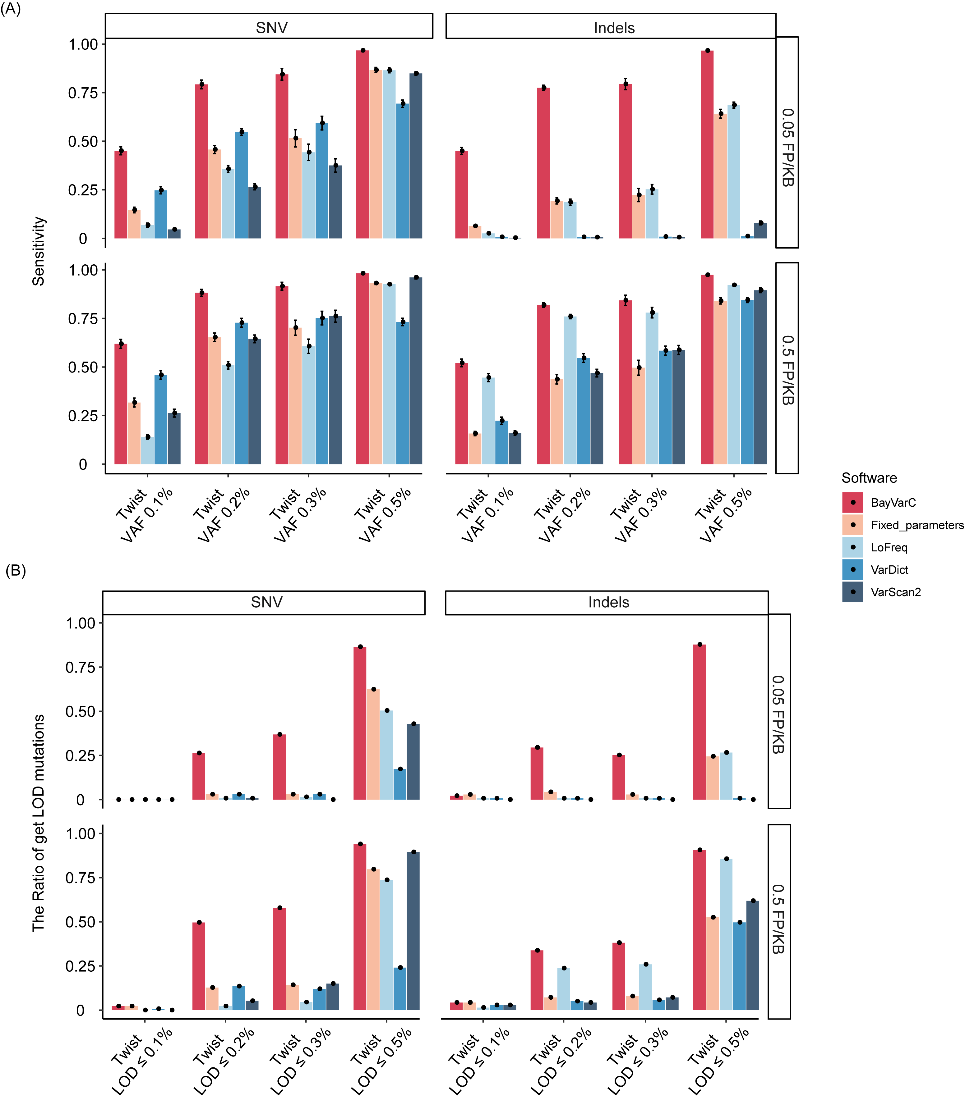

**Supplementary Figure 4:** In-vitro performance of each variant callers on Nova platform. (A) Sensitivity benchmark across variant callers when FPR was fixed at 0.05, 0.5 PF/Kb. (B) LoD benchmark across variant callers when FPR was fixed at 0.05,0.5 PF/Kb.
